# Organism-Environment Topological Interfaces Drive the Origination of Organismal Form

**DOI:** 10.64898/2026.04.05.716476

**Authors:** Weiqi Li, Xudong Zhang

## Abstract

The origin of organismal forms is one of the most enduring unresolved challenges in evolutionary biology. While Darwinian theory and the Modern Synthesis explain adaptive evolution, they do not adequately account for the initial generation of core morphological architectures and rapid diversification events such as the Cambrian explosion. Here we establish that topological interfaces between organisms and their environment are the primary drivers of form origination. Starting from a spherical interface, topological transformations into closed disk or closed cylinder interfaces, governed by resource transport constraints and topological selection, determine the foundational forms of an organism, including shape, size, and complexity. This is supported by our simulations of homeostatic regulation under environmental stoichiometric fluctuations and geometrically allowed morphospace. Furthermore, closed cylinder interfaces generate a directional stoichiometric gradient, which provides a driving force for motion relative to the environment. Validated by empirical data cross kingdom and phylum, our theory yields qualitative and quantitative predictions for key morphological traits, including the species abundance distributions, origination of motility, body size scaling, and explosive diversification, which overcomes long-standing limitations of classical frameworks. This topological paradigm thus provides a unifying mechanism for the origin of biological form.

## 1. Introduction

Morphological evolution is a central theme in biology, crucial for comprehending adaptation, tracing phylogenetic relationships, and unraveling developmental biology^1^. It encompasses the origination and diversification of organismal forms, focusing respectively on how these forms first emerged and how they subsequently diversified and specialized^1^. However, explaining both the origin of organismal forms (e.g., their shape, size, and complexity) and the patterns of their origination (e.g., gradual or burst-like) is a significant challenge in evolutionary theory from Darwinian to the Modern Synthesis^1,2^. These theories are considered to lack a generative mechanism for form generation and can interpret and/or predict only what form will be maintained but not the one that will emerge^3^. The origination of organismal forms thus remains a missing piece in the puzzle of evolutionary theory.

Organisms function as open thermodynamic systems, engaging in exchanges of energy and matter with their surroundings. This interaction enables them to acquire the negative entropy essential for maintaining life^4^. The self-organization theory posits that, under far-from-equilibrium conditions, an open system has the potential to spontaneously progress towards the creation of order through self-organizing processes^5,6^. Within an organismal system, this order-directed process is regarded as evolution, and as such, is expected to give rise to morphological structures^6-10^. There are several straightforward examples at the cellular ^7^, individual, and colony levels that illustrate the formation of biological patterns through self-organization. For instance, the nest-building behavior of fungus-growing *Macrotermes* results in a highly organized structure^6^. However, considering that self-organizing events occur within complex contexts, the currently understood simple mechanisms of self-organization have limited predictive power regarding both the processes and outcomes. Consequently, they are unable to elucidate how organismal forms originated in the real world^6,11^.

Homeostasis is another essential characteristic of organismal forms, and is indispensable for all life-related entities ranging from micelles to the human body^12-14^. Typically, homeostasis entails negative-feedback regulation between the internal and external conditions of complex systems. This mechanism enables organismal systems to maintain a stable chemical composition under environmental fluctuations. Through homeostasis, these systems can stay within viable states regardless of alterations in environmental conditions^15^. Such regulatory processes can be denoted by the coefficient *H* in the formula: *y* = *cx*^1/H^ (where *x, y*, and *c* symbolize resource stoichiometry, organismal stoichiometry, and a constant respectively; *H* ≥ 0)^16^. Furthermore, an organism may exhibit heterogeneous chemical composition or stoichiometric gradients within its body or across distinct tissues and organs. These regions may, in turn, be subject to differential homeostatic regulation^12,17^. Importantly, organisms with high homeostatic capabilities (i.e., large *H* values) have a strong competitive advantage in growth and/or survival under environmental changes^15,16,18^.

This study aimed to propose a theory for the origination of organismal forms. We presented topological models of the interface between organismal systems and their environments, which were formulated into mathematical models based on homeostatic regulation and organismal complexity. Our numerical simulations of competitive advantage and geometric analysis of systemic morphospace support a homeostasis-favored and topology-allowed mechanism for form origination. This mechanism generates key organismal traits, including basic size, shape, complexity, and the driving force of motion. Qualitative and quantitative assessments of these morphospace features match both real-world observations and empirical data at the kingdom and phylum levels, particularly aligning with patterns of animal appearance during the Cambrian period. Thus, we propose a topology-based theory for form origination, whose validity is supported by the foregoing assessments. Nevertheless, we discuss the limitations of this theory in explaining evolutionary dynamic processes, as well as the evolutionary roles of topological interfaces in life’s diversification.

## 2. Model and the Theory

### 2.1 Formulation of the fluctuation in environmental and systemic stoichiometry

In general, organisms obtain energy and matter through the metabolism of ingested resources. Consequently, the exchange of energy and matter between organismal systems (hereinafter referred to as “systems”) and the environment can be simplified to resource intake, as shown in Fig. 1a and b. In our formulation, we adopted a stoichiometric approach to characterize resource intake (Fig. 1b). We denoted the stoichiometry in the environment and within the system as *X*_0_ and *X*_1_, respectively. Thus,the *y* = *cx*^1/*H*^ formula alluded to earlier was rewritten as Eq. (1):

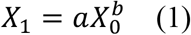

where *a* is a constant, and *b* is the homeostatic regulation constant (i.e., the reciprocal of *H*, with 0 ≤ *b* ≤ 1). The value of *X*_0_ is invariably > 0 and, as the environment changes, it continuously fluctuates around the optimal systemic requirement. Based on this characteristic, we hypothesize that *X*_0_ can be modeled as a sine function of time (*t*). This relationship is briefly presented by the simplified Eq. (2) (Fig. 1c):

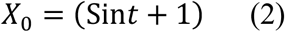

**Fig. 1.**
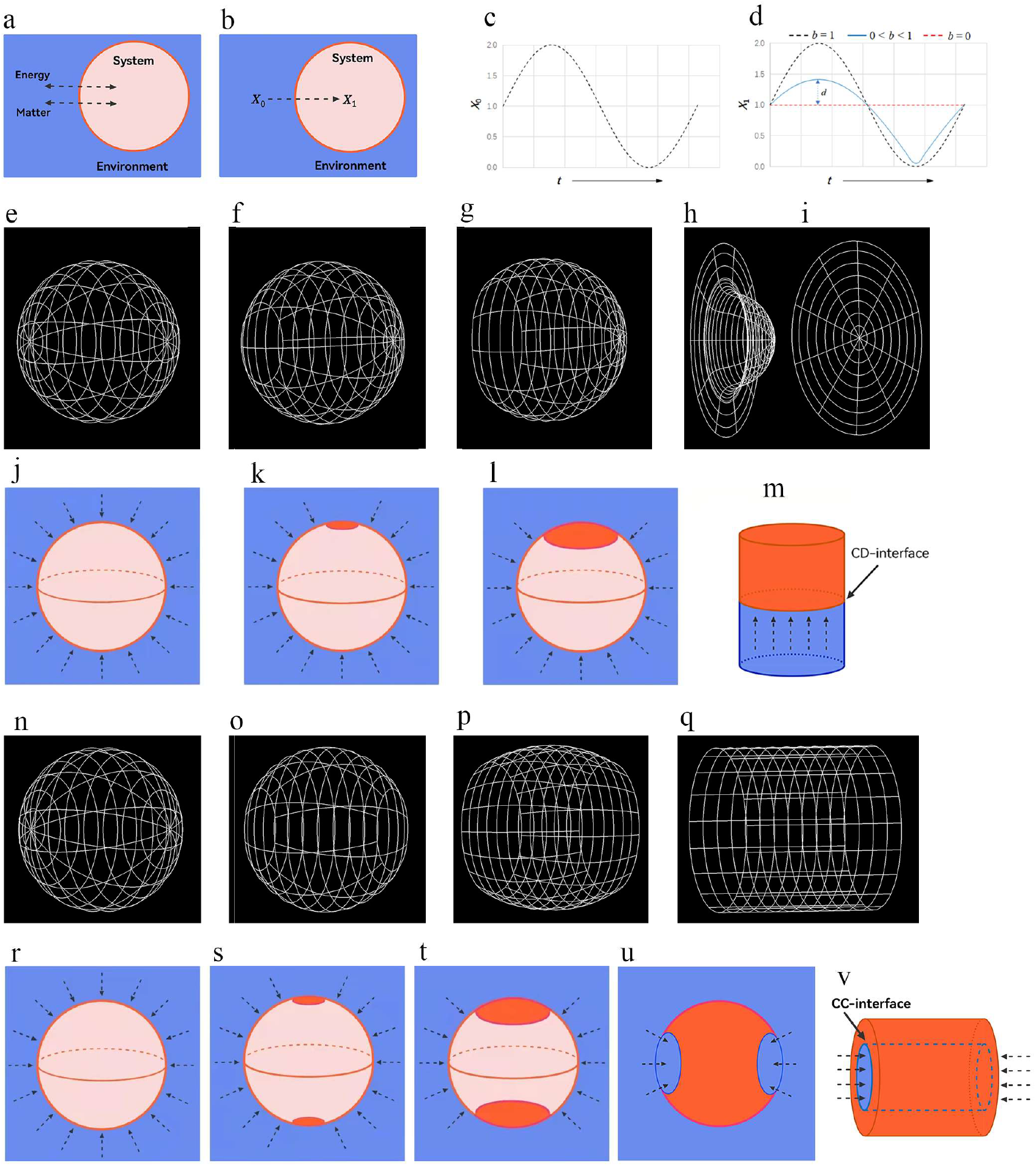
Modeling and illustration of topological interfaces between a system and the environment. (a) Diagram of the thermodynamic model of the system and environment. (b) Diagram of environmental stoichiometry (*X*_0_) and systemic stoichiometry (*X*_1_). (c) Simulation of *X*_0_ fluctuation (Eq. [2]), where *t* is time. (d) Simulation of *X*_1_ fluctuation (Eq. [3]), where *b* is the homeostatic regulation constant, *d* is the deviation of *X*_1_ from *X*_b=ο_ (Eq. [4]), and *t* is time. (e–i) Removing one open disk from a sphere (retaining the boundary) yields a space homeomorphic to a closed disk (CD) upon compactification. (j–m) Appearance of a no-resource-transport (NRT) spot on a spherical interface driving the interface to transform into a CD interface. (n–q) Removing two disjoint open disks and adding boundaries yields a space homeomorphic to a closed cylinder (CC). (r–v) Appearance of two NRT spots on a spherical interface causes the interface to transform into a CC interface. (r–v) Appearance of two NRT spots on a spherical interface, triggering its transformation into a CC interface. **Notes:** Blue area, environment; pink/red area, system; black dotted arrows, direction of energy and/or matter (stoichiometry) flux; red spots, NRT spot formation.

By substituting Eq. (2) into Eq. (1) and performing mathematical simplification (Methods 1 and 2, Fig. 1d), we obtained the expression for the systemic stoichiometry, i.e., Eq. (3):

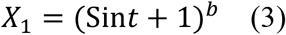

When *b* = 0, the homeostatic regulation of the system reaches its maximum, *X*_1_ = 1 (denoted as *X*_b=ο_ = 1). This means that the systemic stoichiometry remains constant and does not respond to changes in the environmental stoichiometry (Fig. 1d, red dashed line). When *b* = 1, there is no homeostatic regulation in the system, and its stoichiometry changes completely with variations in the environmental stoichiometry (Fig. 1c; Fig. 1d, black dashed line). When 0 < *b* < 1, the fluctuation range of the systemic stoichiometry narrows (Fig. 1d, blue solid line). The deviation of the systemic stoichiometry from that when *b* = 0 is expressed by Eq. (4):

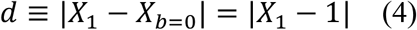

Therefore, we can simulate changes in *X*_1_ to observe the values of *d* under different conditions (Fig. 1d). According to the homeostasis principle^16^, a smaller *d* value indicates a stronger homeostatic regulation ability, which in turn implies a more significant competitive advantage. Thereafter, we use *d* to distinguish the level of survival capacity of a system.

### 2.2 Three topological structures of system-environment interfaces

All modern cellular life evolved from the unicellular last universal common ancestor (LUCA)^19,20^. Unicellular systems absorb resources isotropically and radially from the environment. This gives rise to a **sp**herical system–environment interface (SP, Fig. 1e and j). We propose this spherical interface as the origin of morphological diversification. From a topological perspective, a spherical interface must contain at least one tangential singular region, as required by the Hairy Ball Theorem ^21^. Removing a disk-shaped region from this sphere yields a **c**losed-**d**isk (CD) interface (Fig. 1e–i). We therefore hypothesize that the tangential gradient field of surface properties (e.g., permeability, material exchange efficiency) forms a specialized region lacking resource uptake, termed a **n**on-**r**esource **t**ransport (NRT) spot. This transitions the system–environment interface from SP to CD (Fig. 1j–m). Furthermore, the emergence of two tangential singular regions and removal of two disk-shaped regions from the sphere generate a **c**losed-**c**ylinder (CC) interface (Fig. 1n–q). Correspondingly, we hypothesize that two NRT spots arise on the system–environment surface (Fig. 1r–u), converting the interface to the CC form (Fig. 1v). We thus hypothesize that the system–environment interface adopts three distinct topological configurations—SP, CD, or CC—that underlie all primary morphological structures.

### 2.3 Homeostatic and geometric constraint on complexity development in a system with CD interface

We then tested whether these three interface types can define the fundamental forms of organisms by examining their constraining effects on two key aspects: **h**omeostatically **f**avored (HF) and **g**eometrically **a**llowed (GA) development of systemic complexity. For the HF analysis of each interface type, we set up three systems comprising single, double, and triple stoichiometric subsystems respectively to represent increasing systemic complexity (Fig. 2), and calculated the changes in systemic *d* values (Eq. [4]) in response to increasing systemic complexity. Systems with lower *d* values tend to enhance their complexity via their topological interfaces, thereby enhancing their survival capacity. For the GA analysis of each interface type, we analyzed the spatial expansion of subsystems along the interface’s normal direction from a geometric perspective, and thus inferred the allowed morphospace for systemic development. By combining these two aspects, we can obtain the fundamental forms of the system under each topological interface type.

**Fig. 2.**
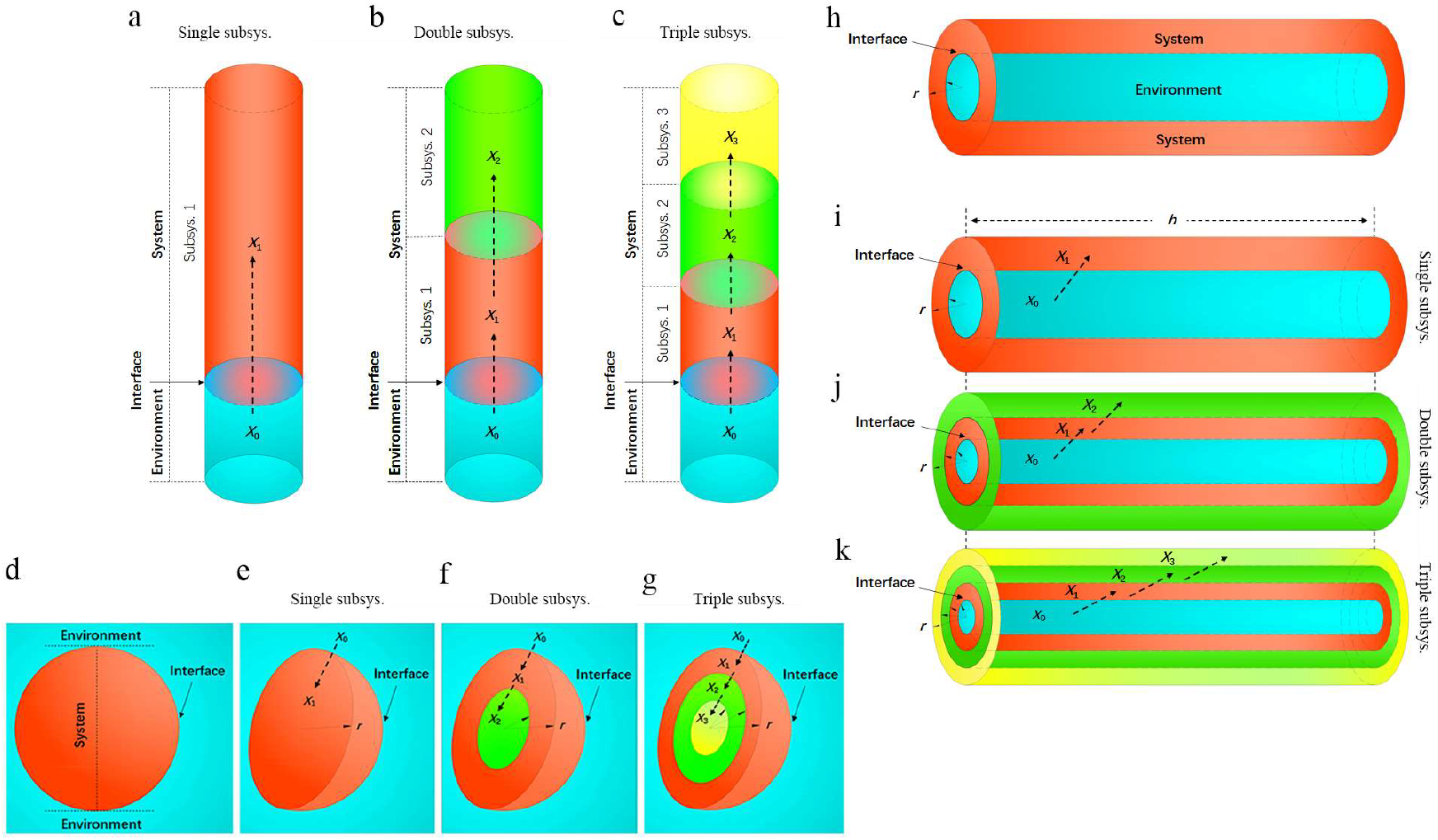
Illustration of systemic complexity models. Diagrammatic representations of (a–c) models of closed disk (CD)-interface systems comprising single, double, and triple subsystems; (d) sphere (SP)-interface system; (e–g) spherical-interface systems comprising single, double, and triple subsystems; (h) closed-cylinder interface system; (i–k) closed cylinder (CC)-interface systems comprising single, double, and triple subsystems. **Notes:** Blue, environment; red, system/subsystem 1; green, subsystem 2; yellow, subsystem 3; black dotted arrows, direction of stoichiometry flux.

We first examined the CD-interface system using the above method. Firstly, we investigated the homeostatic tendency by setting up systems comprising single (*X*_S_), double (*X*_D_), and triple subsystems (*X*_T_), whose stoichiometries can be expressed by Eqs. (5), (6), and (7) respectively (Fig. 2a−c, Method 3):

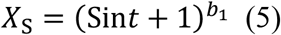

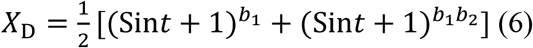

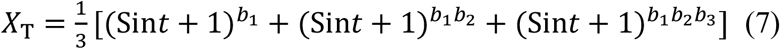

where *b*_1_, *b*_2_, and *b*_3_ are the homeostatic regulation constants of subsystems 1, 2, and 3, respectively. Simulated calculations of the three systemic stoichiometry (top panel of Fig. 3) showed that a system with more subsystems exhibits a lower *d* value (with *X* closer to 1). This indicates that CD-interface systems tend to increase their complexity. Secondly, we investigated the space allowed for subsystem development by the CD interface. The spatial constraint of the CD interface on the system can be determined by its topological characteristics (Fig. 2a–c). The system obtains resources from one side of the interface, which means that subsystems can only develop toward the other side of the interface, and the volume of the expanded space is linearly correlated with the height along the normal direction of the interface. Therefore, the spatial expansion of the CD-interface system is not limited, and the spatial shape is not predetermined.

**Fig. 3.**
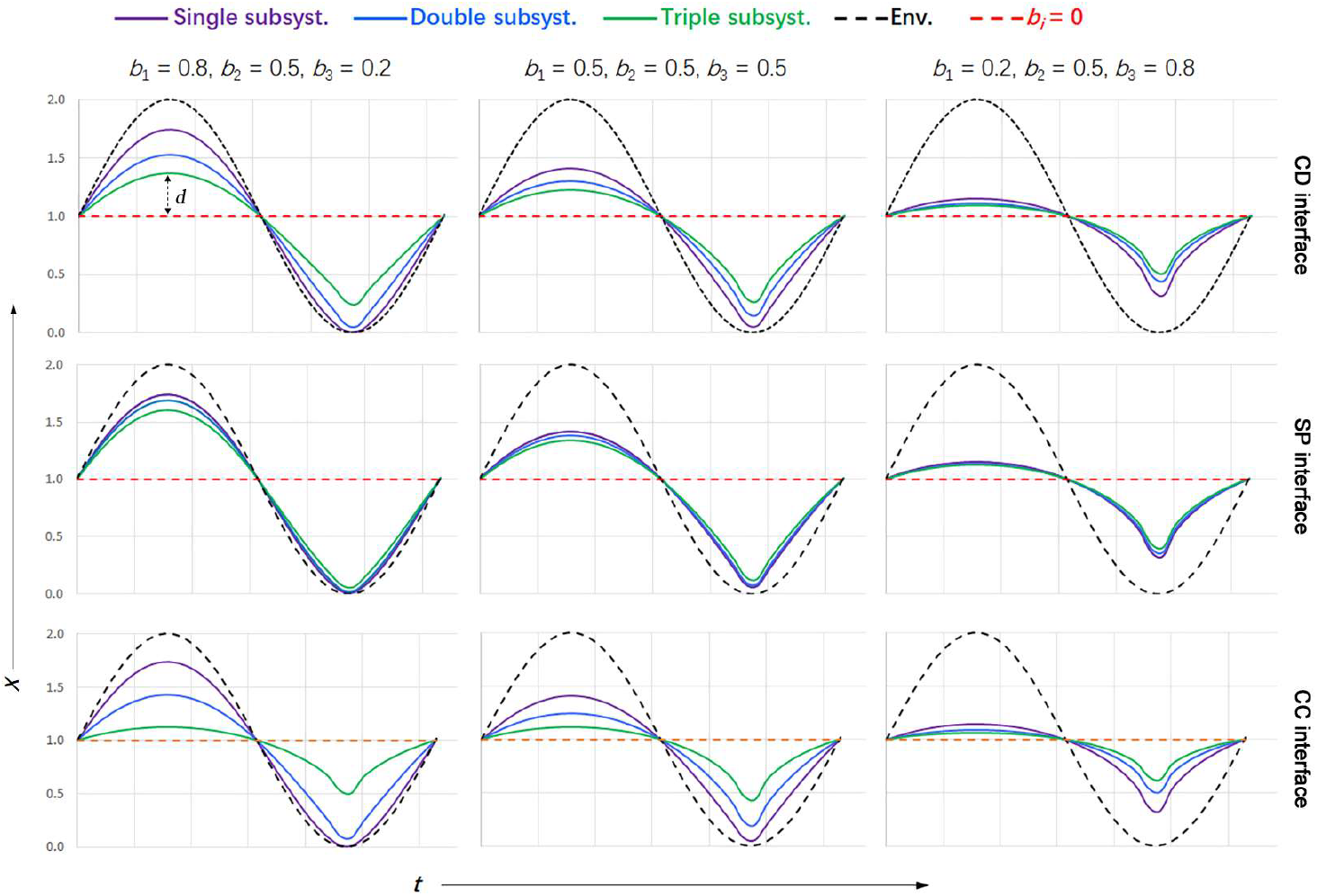
Simulated systemic stoichiometry with increasing complexity under different homeostatic regulation levels. Definitions: *X*, stoichiometry; *t*, time; *d* is the deviation of *X*_*i*_ from *X*_b=ο_. **Notes:** Solid lines, stoichiometry of systems comprising single (purple), double (blue), or triple (green) subsystems; black dotted lines, stoichiometry of environment (equivalent to that of a system with homeostatic regulation constant *b* = 1); red dotted lines, stoichiometry of a system with a homeostatic regulation constant *b*_*i*_ = 0 (*i* = 1, 2, or 3); *b*_1_, *b*_2_, and *b*_3_, homeostatic regulation constants of subsystems 1, 2, and 3, respectively (Methods 3–5). CD, closed disk; SP, sphere; CC, closed cylinder.

Taken together, the fundamental form of a system with a CD interface tends to develop complex subsystems in one direction of the interface (Fig. 2a−c).

### 2.4 Homeostatic and topological constraint on complexity development in a system with SP interface

We examined the HF and GA form for the system development of SP-interface systems. In terms of the system’s HF form, the stoichiometry of systems comprising single (*X*_S_), double (*X*_D_), and triple (*X*_T_) subsystems (Fig. 2d) can be expressed by Eqs. (8), (9), and (10) respectively (Fig. 2e–g, Method 4 and Table S1):

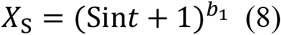

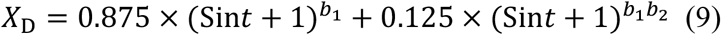

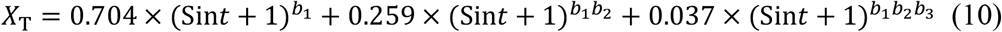

Simulated calculations of systemic stoichiometry (Fig. 3, middle panel) showed that systems with more or fewer subsystems exhibit similar *d* values (i.e., the distance between *X* and 1 is comparable across these systems), which means that their homeostatic potential is similar regardless of the number of subsystems. This result indicates that the complexity of SP-interface systems has no competitive advantage.

In terms of the GA form for system development, the system can obtain resources from the outside through the interface, so subsystems can only develop inward along the normal direction of the interface (Fig. 2e–g). Assuming the sphere volume is 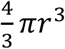, the results showed that as the radius (*r*) decreases along the normal direction, the space for subsystem development decreases with the cube of *r*.

Taken together, systems with an SP interface exhibit minimal tendency and limited space for subsystem development, and are thus inferior to those with a CD interface in both aspects.

### 2.5 Homeostatic and topological constraint on complexity development in a system with CC-interface system

For systems with a CC-interface, in terms of the system’s HF form, the stoichiometry of systems comprising single (*X*_S_), double (*X*_D_), and triple subsystems (*X*_T_) can be expressed by Eqs. (11), (12), and (13) respectively (Fig. 2i–k, Method 5 and Table S2):

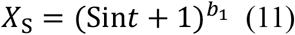

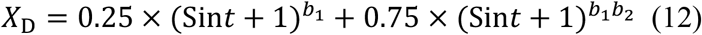

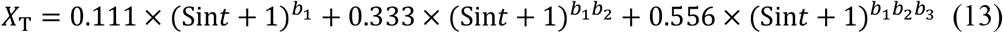

Simulated calculations of systemic stoichiometry (Fig. 3, bottom panel) showed that systems with more subsystems exhibit lower *d* values (with *X* closer to 1); and, under the same *b*_i_ conditions, the negative correlation between complexity and *d* values in CC-interface systems is much more significant than in CD-interface ones (Fig. 3, top panel). In other words, the increased complexity of CC-interface systems enhances their competitive advantage.

In terms of GA form, the system acquires resources from the inside of the interface, so subsystems can only develop outward from the outside of the interface along the normal direction of the interface (Fig. 2i–k). Assuming the cylinder volume is *πr*^2^*h*, the development space for subsystems will expand quadratically as the radius (*r*) increases along the normal direction. Therefore, relative to CD-interface systems, CC-interface systems offer greater space for subsystem development.

Taken together, systems with a CC interface have a strong tendency and ample space to develop subsystems.

### 2.6 Relative motion tendency of entrance-exit directional resources in the CC-interface system

We further investigated the additional effect of the CC-interface on organismal form. Specifically, when environmental resource availability is uneven, leading to resource intake via a single port (entrance, Fig. 4a), this unevenness results in a continuous decrease in stoichiometry along the system’s cylindrical direction, as well as the spontaneous formation of stoichiometric gradients both perpendicular to the interface (along the normal direction) and parallel to the interface (Fig. 4a). The gradient perpendicular to the interface (along the normal direction) also exists in SP- and CD-interface systems (Fig. 1j and m), while the gradient parallel to the interface is unique to CC-interface systems (Fig. 4a). We thus investigated whether parallel gradients would generate special morphological features. We posit the existence of three systems parallel to the CC interface, which contain one, two, and three subsystems, respectively (Fig. 4b-d). When environmental stoichiometric flux passes through the cylinder, these three systems generate zero gradient, a single gradient, or two stoichiometric gradients, respectively. The stoichiometry of these three systems can be expressed by Eqs. (14), (15), and (16) respectively (Method 6):

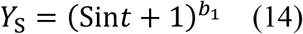

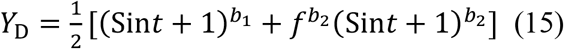

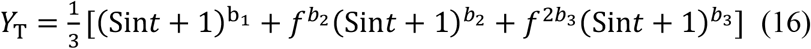

where *f* is the gradient coefficient of the subsystem, 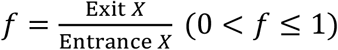. The numerical simulation of *Y*_D_ (Eq. 15 and Fig. S1) and the corresponding deviation *d*_D_ showed that the stoichiometric gradient affects system homeostasis (*d* decreases as *f* decreases); moreover, when subsystems are arranged along the CC interface with *b*_1_ < *b*_2_, their homeostatic regulation ability is stronger than when *b*_1_ = *b*_2_ or *b*_1_ > *b*_2_ (Fig. 5). The simulation of *Y*_T_ (Eq. 16 and Fig. S2) yielded the same results.

**Fig. 4.**
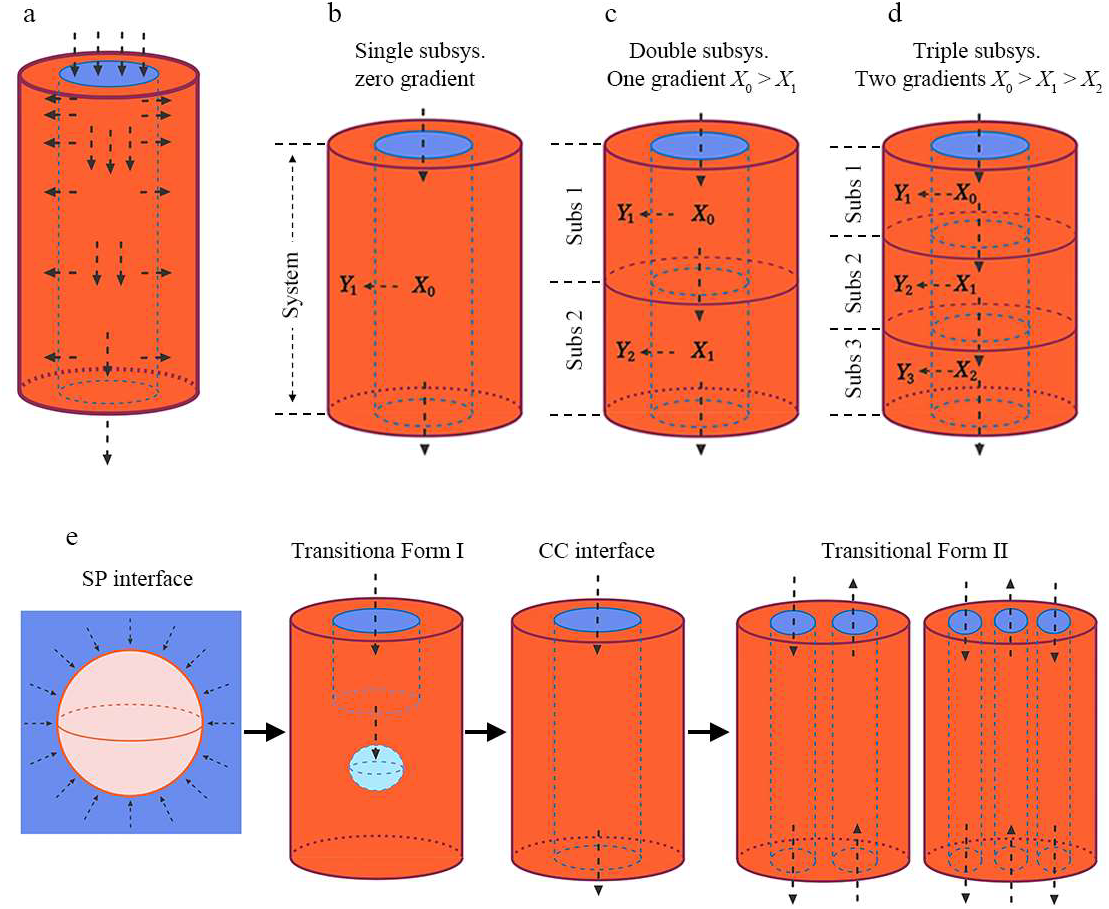
Stoichiometric gradients in closed-cylinder (CC) interface system and transitional forms (TF-I and TF-II). (a) Two orthogonal directions of stoichiometry flux; one normal to the interface, and the other parallel to interface (along the environmental cylinder). Diagrams of systems comprising (b) a single subsystem (no stoichiometry gradient along the environmental cylinder); (c) two subsystems (one stoichiometry gradient, *X*_0_ > *X*_1_); (d) three subsystems (two stoichiometry gradients, *X*_0_ > *X*_1_ and *X*_1_ > *X*_2_). (e) Transitional sequence from spherical (SP), closed cylinder (CC), TF-I (half-penetrated environmental cylinder with cavity + vacuole), to TF-II (multi-penetrated environmental cylinders). **Notes:** Blue area, environment; red area, system; black dotted arrow, direction of resource flux; dotted blue sphere, resource vacuole; *X*_0_, *X*_1_, and *X*_2_, environmental stoichiometry; *Y*_1_, *Y*_1_, and *Y*_2_, subsystemic stoichiometry.

**Fig. 5.**
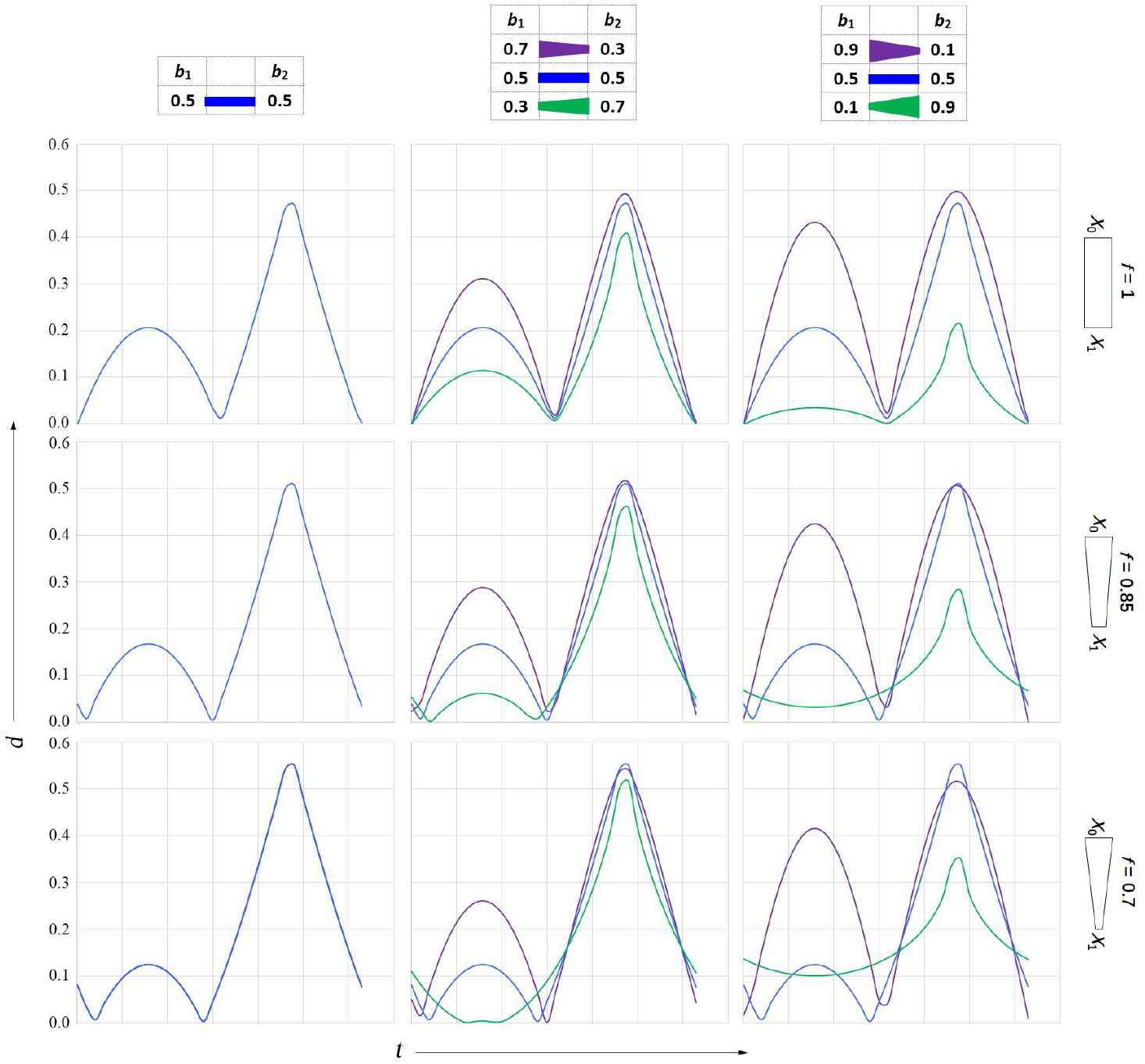
Simulation of deviation 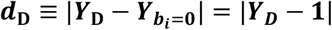 (*i* = 1 or 2, Eq. [4]). **Definitions:** *d*, deviation of systemic stoichiometry from that at *b* = 0; *t*, time; *b*_1_ and *b*_2_, homeostatic regulation constants of subsystems 1 and 2, respectively. **Notes:** Purple, blue, and green lines denote *d* values for systems with subsystemic *b*_1_ >, *b*_1_ = *b*_2_, and *b*_1_ < *b*_2_, respectively; gradient coefficient, 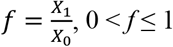 (Fig. 5c).

These results indicate that once resources enter the cylinder and are taken up by the system, a stoichiometric direction favored by system homeostasis is formed, specifically weaker homeostatic regulation at the entrance and stronger at the exit. Consequently, the gradient parallel to the interface is sustained, thus continuously driving relative movement between the resources and the system.

### 2.7 The transitional forms of topological interfaces between system and environment

In principle, other than the SP, CD, and CC interfaces, the existence of additional interfaces cannot be ruled out. Specifically, from a geometric perspective, such interfaces may exist in two possible forms. The first is a form transitioning from the SP interface to the CC interface (Fig. 4e, referred to as **T**ransitional **F**orm I, TF-I), which can be regarded as having “half”-penetrated environmental cylinders. Its tendency for spatial expansion and stoichiometric gradients parallel to the cylinders are, respectively, in a transitional state between low/absent (close to the SP interface) and high/present (close to the CC interface). The second is a form of the CC interface evolving toward greater complexity (Fig. 4e, referred to as **T**ransitional **F**orm II, TF-II), which has multiple environmental cylinders penetrating the system. For this form, since each environmental cylinder generates its own stoichiometric direction (Fig. 4e), their different directions will counteract each other, thereby making the homeostasis-favoring property of the TF-II system weaker than that of the CC-interface system. In summary, TF-I and -II may exist, but their homeostatic tendencies are weaker than those of systems with the SP or CC interface.

### 2.8 A model for the origination of organismal forms

The modeling presented above establishes a generative mechanism for the origination of organismal forms. Mathematical simulation of stoichiometric homeostasis and geometrical analysis demonstrate the mechanism to be homeostatically favored and geometrically allowed. On this basis, we propose a theory for the origination of organismal forms:

1. Organismal forms originate from the SP interface formed between a unicellular/aggregated cellular system and the environment, and this interface transforms into a CD interface or a CC interface (Fig. 2). These three types of topological interfaces define the main morphological features of organismal systems: shape, size, and complexity.
2. The SP-interface system has a morphospace constrained inward cubically and does not favor complexity; The CD-interface system develops linearly from one side of the interface with an indefinite shape, favoring complexity; The CC-interface system exists in the form of an environmental cylinder penetrating the systemic body, with a morphospace expanding outward quadratically, and strongly favors complexity (Fig. 2).
3. The CC-interface system generates a stoichiometric gradient with the entrance-exit direction parallel to the cylinder and can drive relative movement between the system and the environment (Fig. 4a–d).
4. There are two types of transitional form systems (TF-I and -II), with half-penetrated and multi-penetrated environmental cylinders, respectively. They comprise some features of both the SP-interface system and the CC-interface system, and their homeostatic tendencies are weaker than those of the latter two (Fig. 4e).

## 3. Predictions and Verifications

We verified the validity of our theory by assessing its ability to predict organismal forms and their natural occurrence frequency (Fig. 6a). Specifically, we analyzed the resource-intake structure of organisms corresponding to each topological interface type, deduced their predicted morphological features, counted such organisms in natural systems, and further quantified the match between predicted and observed features as well as the quantities of these organisms.

**Fig. 6.**
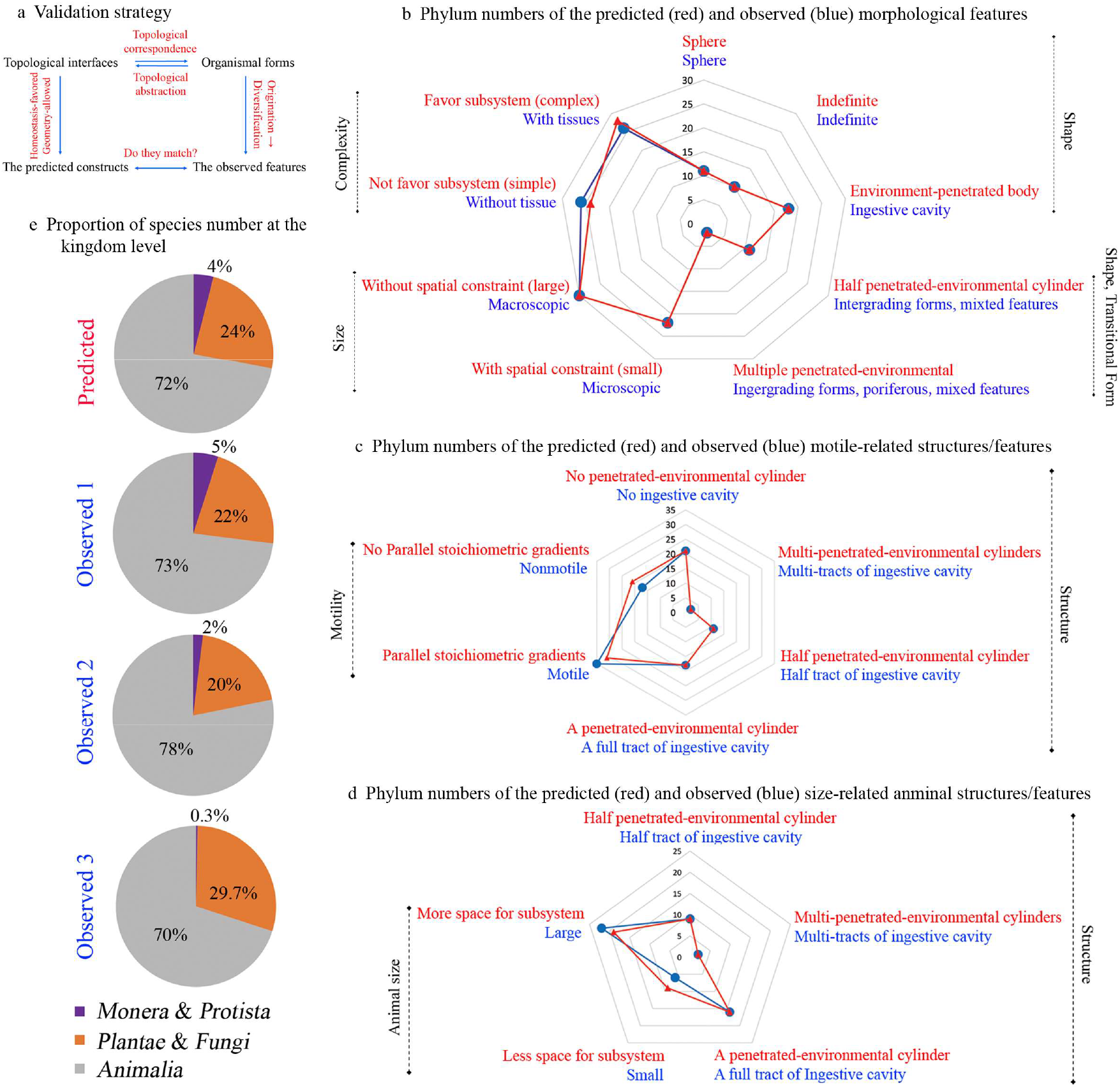
Comparison of predicted and observed morphological forms. (a) Schematic of the validation strategy. (b) Numbers of phyla exhibiting indicated morphological features. Nine categories include three shape-related, two shape and transitional form-related features, two size-related, and two complexity-related (details in Data S2). (c) Numbers of phyla with indicated structures and motile features. Six categories include one structure without parallel stoichiometric gradient, three structures with parallel stoichiometric gradient, and two motile features (details in Data S3). (d) Numbers of phyla with indicated structures and relative animal size. Five categories include one structure providing limited subsystem space, two structures providing ample subsystem space, and two size features (details in Data S4). (e) Kingdom-level species proportion. Predicted proportions correspond to those in Fig. S3; observed proportions are from previous studies (Table S5). Topological interface assignments for each kingdom are provided in Table S3.

### 3.1 All organismal forms at the kingdom and phylum levels can be represented by the three topological interfaces and two transitional forms

If our theory can account for the origin of organismal forms, then all organismal forms should be representable by the SP, CD, and CC interfaces, together with the TF-I and TF-II transitional forms (Fig. 6a). We first test whether this claim holds true.

As understanding of organismal morphology and evolution has advanced, organisms have historically been classified into two, three, four, and five kingdoms ^22,23^ (Table S3).

- Initially, organisms were categorized into the *Plantae* and *Animalia* kingdoms, primarily based on their resource-intake structures. Specifically, nutrient absorption via roots (for *Plantae*) and food ingestion through the gut cavity (for *Animalia*), respectively ^23^. These two structures can be explained by the CD and CC interfaces, respectively: the former enables resource-intake from one side of the interface, while the latter from the inside of the interface (Fig. 2a, 2i, and Table S3), respectively.
- Following the discovery of microorganisms, organisms were further classified into the *Protista, Plantae*, and *Animalia* kingdoms^22,23^. *Protista* (unicellular and nutrient-absorptive) can be accounted for by the SP interface, which allows resource intake from the outside of the interface (Fig. 2 and Table S3).
- The widely accepted five-kingdom classification also relies on these same nutritive modes ^22,23^. Specifically, the kingdoms of *Monera* and *Protista* (unicellular, absorptive), *Plantae* and *Fungi* (multicellular, absorptive), and *Animalia* (multicellular, ingestive) can each be explained by the SP, CD, and CC interfaces, respectively, with *Monera* and *Protista* corresponding to the SP interface, *Plantae* and *Fungi* to the CD interface, and *Animalia* to the CC interface (Fig. 2 and Table S3).

We further assessed resource-intake structures across organisms at the phylum level^22,23^, finding that 39 out of 52 phyla can be represented by SP-, CD-, or CC-interface systems (Data S1). The remaining 13 phyla, however, possess ingestive cavities that diverge into two groups with distinct structures:

- The first group, consisting of 11 phyla, has partial cavity tracts (e.g., via endocytosis or cytostome-vacuoles) and is hypothesized to be transitional forms during the evolutionary transition from absorption to ingestion (i.e., internalization)^22^. These phyla can be accounted for by TF-I with half-penetrated environmental cylinders (Fig. 4e and Data S1).
- The second group (2 phyla: *Porifera* and *Archaeocyatha*) possesses multiple cavity tracts and ingests material via oscula. This group is speculated to be either transitional forms^24^ or a side branch^22^ during animal evolution and can be explained by TF-II with multi-penetrated environmental cylinders (Fig. 4e and Data S1).

Taken together, the SP, CD, and CC interfaces are capable of representing all organismal forms at the kingdom level, while all organismal forms at the phylum level can be represented by the SP, CD, and CC interfaces, plus the two transitional forms (TF-I and TF-II). The results validated that the predicted and observed quantities of organisms with these forms are aligned.

### 3.2 Predictions of the fundamental shape, size, and complexity of organisms

Based on the homeostatically favored and geometrically allowed constructs of SP, CD, CC interfaces, and TF-I and TF-II forms (Fig. 2, Fig. 4e, and Data S1), we predicted the fundamental morphological features of organisms across 52 phyla and classified these features into 9 categories for ease of description (Fig. 6b and Data S2):

1. Shape-related features (3 categories): The SP, CD, and CC interfaces generate spherical, indefinite, and environment-penetrated body forms in organisms, respectively.
2. Size-related features (2 categories): The SP interface imposes spatial constraints on subsystem development, resulting in small organisms; in contrast, the CD and CC interfaces provide space for subsystem development, leading to large organisms.
3. Complexity-related features (2 categories): The SP interface does not favor subsystem development, generate simple organisms; whereas the CD and CC interfaces favor such development, enabling the formation of complex organisms.
4. Transitional form-related features (2 categories): TF-I provides a certain degree of homeostasis and constraints for subsystem development between the SP and CC interfaces, thus generating organisms with morphological features combining those of both interfaces; TF-II offers tendencies for multi-directional subsystem development, thereby generating diverse organisms with weaker homeostasis than that of CC-interface systems.

We then compared the numbers of phyla with predicted morphological features and those with corresponding observed features, and found that they exhibit strong consistency (Fig. 6b, Data S2, and Table S4). For example: 22 and 30 phyla were predicted to be small and large organisms, respectively, while 22 and 30 phyla were observed to be microscopic and macroscopic organisms, respectively; 24 and 28 phyla were predicted to be simple and complex systems, respectively, whereas 26 and 26 phyla were observed to be organisms without tissues and those with tissues, respectively; 11 and 2 phyla were predicted to be TF-I and TF-II transitional forms, respectively, and these phyla were observed to be transitional groups, typically *Mesozoa* and *Porifera*. These organisms lack tissues but exhibit macroscopic forms, i.e., they combine the predicted features of the SP and CC interfaces. Taken together, the results provide direct support for the validity of our theory.

### 3.3 Prediction of organisms capable of generating relative motion with their environment

Our theory predicts the following: Organisms with CC-interfaces, TF-I, and TF-II structures possess environment-penetrated cylindrical structures, within which there exist directional resource stoichiometric gradients parallel to the cylinders. These gradients can drive relative motion between resources and the organismal system; thus, such organisms develop a tendency toward motility relative to the environment and are classified as animals (Fig. 4). In contrast, organisms with SP and CD interfaces lack such structures, cannot develop a tendency for relative motion with the environment, and therefore do not exhibit motility. Among the 34 phyla observed in the real world to have motile characteristics, 30 (accounting for 88%) were successfully predicted by our theory (Fig. 6c and Data S3). The alignment between predicted and observed results is statistically significant (Table S4), indicating that our theory can predict the generation of relative motion between organisms and the environment.

### 3.4 Prediction of body size scaling in animals at phyla level

Based on the traditional morphological taxonomy of animals, the Kingdom *Animalia* is divided into *Protozoa* and *Metazoa*^25^, encompassing 28 phyla (Data S4). Our theory can further predict that phyla with half-penetrated environmental cylinders (TF-I) will provide less space for subsystem development compared with those with multi-penetrated environmental cylinders (TF-II) or fully penetrated (CC, Fig. 4e). One outcome of this prediction is that the size of TF-I organisms should be smaller than that of TF-II or CC-interface organisms. This prediction was verified by observations that protozoans are usually smaller than metazoans^26,27^ (Fig. 6d, Table S4, and Data S4). This suggests that our theory can predict the relative size of animals.

### 3.5 Prediction of species abundance distributions at kingdom level

Our theory can predict species abundance distributions (species proportion) at the kingdom level by assuming that a system develops subsystems to yield diversity and that the morphospace volume of newly developed subsystems is proportional to the number of newly created species (see details in Fig. S3). Qualitatively, the tendency of expansion of subsystem morphospace increases as follows: the SP < the CD < the CC interface (Figs. 2 and 3). Thus, diversity should be the lowest in SP- and highest in CC-interface organisms. Considering that at the kingdom level, SP-interface organisms correspond to *Monera* and *Protoctista*, CD-interface organisms correspond to *Plantae* plus *Fungi*, and CC-interface organisms correspond to Animalia, species numbers can thus be predicted as follows: *Monera* + *Protoctista* < *Plantae* + *Fungi* < *Animalia*. This is in accordance with the empirical data (Table S5)^28,29^.

Quantitatively, the predicted proportions of species numbers among SP-, CD-, and CC-interface organisms were as follows: 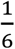 : 1: 3 = 4.0%: 24.0%: 72.0% (see details in Fig. S3). Surprisingly, this matches the observed proportions of species numbers: 5% in *Monera* and *Protoctista*, 22% in *Plantae* and *Fungi*, and 70%–73% in *Animalia*^28^ (Fig. 6e and Table S5). Particularly, our predicted species-number ratio of 1:3 between CD- and CC-interface organisms is exactly equal to the ratio between *Plantae* plus *Fungi* and *Animalia* (reported as 1:3.18, 1:3.93, or 1:2.4 by different research groups).

### 3.6 Prediction of the pattern of animal appearance during evolution

The pattern of animal (particularly metazoans) appearance in evolutionary history is a critical issue for explaining the Cambrian explosion, when major animal groups (more than 100 extant phyla and classes of animals) suddenly appeared over a relatively short period of time^30,31^. Rapid morphological diversification is unlikely to result from constant and incremental variation^30,31^. However, using the features of the CC interface (all metazoans are CC-interface organisms, Data S1), we can predict the pattern of animal appearance as follows:

- The formation of the CC-interface system (which requires two NRT spots) is more difficult than that of the CD-interface system (which requires only one NRT spot) (Fig. 2). Therefore, animals should have appeared after plants over the course of evolutionary time.
- The CC-interface system enables quadratic expansion of morphospace and exhibits the strongest tendency toward systemic complexity. In contrast, the SP-interface system exhibits inward cubic constraint on morphospace and does not favor complexity, while the CD-interface system expands morphospace linearly and favors complexity. Therefore, the appearance of animals should have been the most rapid and have shown the greatest diversity.

These qualitative predictions can explain the patterns of animal appearance during the Cambrian explosion very well, and thus suggest that the unique pattern of animal appearance is primarily attributed to their CC-interface structure.

## 4. Discussion

Building on the topological structure of the environment-organismal system interface and the environmental stoichiometric role in the system’s homeostasis, we developed a model that accounts for the origin of organismal form. Using this model, we predicted and explained six fundamental issues tightly linked to the origin of organismal morphology: the forms of organisms at the kingdom and phylum levels; the fundamental shape, size and complexity of organisms at the phylum level; the relative sizes of animals at the phylum level; the origination of the driving force for organismal motion relative to environment; the species abundance distributions at the kingdom level; and the pattern of animal appearance during evolution that can account for the Cambrian explosion.

Three points are noteworthy: First, all of these predictions and explanations align closely with empirical data and real-world observations; Second, these predictions are all endogenously derived from our model, suggesting that the model is a generative one; Third, these predictions range from qualitative (e.g., the relative sizes of animals, the origin of the driving force for motion) to semi-quantitative (e.g., the morphospace expansion of plants is linear while that of animals is quadratic) and quantitative (e.g., the proportion of species numbers).

### 4.1 The topological role of the system-environment interface may be one of the independent factors in defining form origination

Biological evolution is typically driven by two broad types of factors: systemic and environmental. This classification aligns not only with the shared “variation and natural selection” framework of both Darwinian and Modern Synthesis theories^1,3^ but also with empirical evidence. For instance, systemic factors include genetic variation, genetic drift, sexual selection, developmental constraints, and epigenetics, while environmental factors encompass natural selection, geographical isolation, climate change, and geological events. However, our theory reveals that the system-environment interface defines the primary spatial relationship between these two, and this definition is independent of both factor types. This suggests that alongside systemic and environmental factors, the system-environment interface constitutes a third independent driver of evolution. These three factors act as distinct “players” collaborating to shape life’s evolution. Consequently, many unresolved debates in evolutionary biology may stem from the omission of this third player, the system-environment interface. Examples include disputes over the origin of fundamental organismal forms^1,3^, debates over whether self-organization alone is sufficient to account for biological structure formation^6^, and inadequate explanations for the Cambrian explosion^30,31^.

### 4.2 A Comparative analysis of three evolutionary theories in explaining the Cambrian Explosion

The Cambrian Explosion is one of the most significant events in evolutionary history. It not only profoundly shaped the origin and diversification of organisms (particularly metazoans) but also became critical for testing evolutionary theory. We therefore evaluated our model’s ability to explain features of the Cambrian Explosion. The event exhibits five core features: temporal concentration, morphological complexity leap, rapid diversification, occurrence only in the animal kingdom, and uniqueness (i.e., occurring only once) (Table S6).

As is well known, Darwinian theory struggles to explain the Cambrian Explosion. In fact, it fails to account for any of these five features (Table S6)^1,31,32^. The Modern Synthesis partially explains three features (temporal concentration, morphological complexity leap, and rapid diversification) but cannot explain why the event was restricted to animals or why it was unique (Table S6)^31-33^. In contrast, our model explains all five features by leveraging the overwhelming superiority of subsystem development in CC-interface system over that in SP- and CD-interface systems (Table S6). Importantly, these explanations are generated directly and spontaneously from the model. Furthermore, the CC-interface system burst when its superiority in subsystem development aligned with the habitable environment in Cambrian oceans, such as rising oxygen levels and the removal of dissolved barium ions. This explains another puzzle of the Cambrian Explosion: why it specifically occurred in the Cambrian. These analyses suggest that our model has advantages over the two previous theories in explaining the Cambrian Explosion.

### 4.3 Our model has limitations in explaining evolutionary phenomena related to rate and mechanism

Our model employs homeostatic and geometric factors to account for the origin of organismal morphology during evolution, and is fundamentally grounded in thermodynamic principles. Therefore, its explanatory power is limited when it comes to evolutionary phenomena involving the kinetic factors, such as rate, timing, and molecular mechanism. This can be illustrated from three cases.

- Firstly, our model offers a sequential origination of SP-, CD-, and CC-interface system, meaning that animals appeared after plants. However, when addressing the kinetic question of “how long afterward”, our model provides no insight. Instead, environmental factors might provide the timing: the emergence of a habitable ocean environment during the Cambrian period triggered the animal burst^34,35^.
- Secondly, our model posits that organisms with a CC-interface naturally generate forces driving their movement relative to the environment. This prediction encompasses the animal kingdom; in other words, it holds true for the vast majority of motile organisms. However, there are exceptions: certain unicellular organisms can move using flagella, cilia, or pseudopodia without a CC-interface, and our model fails to explain these cases. The reason may lie in the fact that locomotion evolution in these organisms is governed by molecular mechanism. For example, the complex protein assemblies of the rotary motor in flagellar movement.
- Thirdly, the question of how homeostasis-favored and topology-allowed morphologies are genetically inherited involves DNA-related molecular processes and is therefore a kinetic rather than thermodynamic issue. Our model cannot directly address this mechanism, at least not at present.

### 4.4 Form origination based on topological interfaces could be a distinguishing stage between Chemical Evolution and Biological Evolution

Traditionally, it is widely accepted that life evolution proceeds through two basic stages: Chemical Evolution and Biological Evolution^36^. In the former, inorganic chemicals give rise to biomolecules and membranes. In the latter, lower organisms evolve into higher organisms with remarkable diversity in shape, size, and complexity, which is often vividly depicted as “the tree of life”. Given that the mechanisms underlying morphological origination differ from those underlying diversification^3^, which is supported by our theory, we thus suggest a “middle stage” that exists between chemical and biological evolution. This middle stage is analogous to a “bud” in the transition from a seed to a tree: the bud is not as complex or mature as the tree, but it establishes the primary framework for the tree’s growth. In this sense, our theory likely depicts “the bud of life” (Fig. S4). Most importantly, the constraint shaping life evolution in this middle stage is topological interfaces, a constraint distinct from those acting on chemical and biological evolution in the other two stages. Therefore, the middle stage could be termed “Topological Evolution” (Fig. 7).

**Fig. 7.**
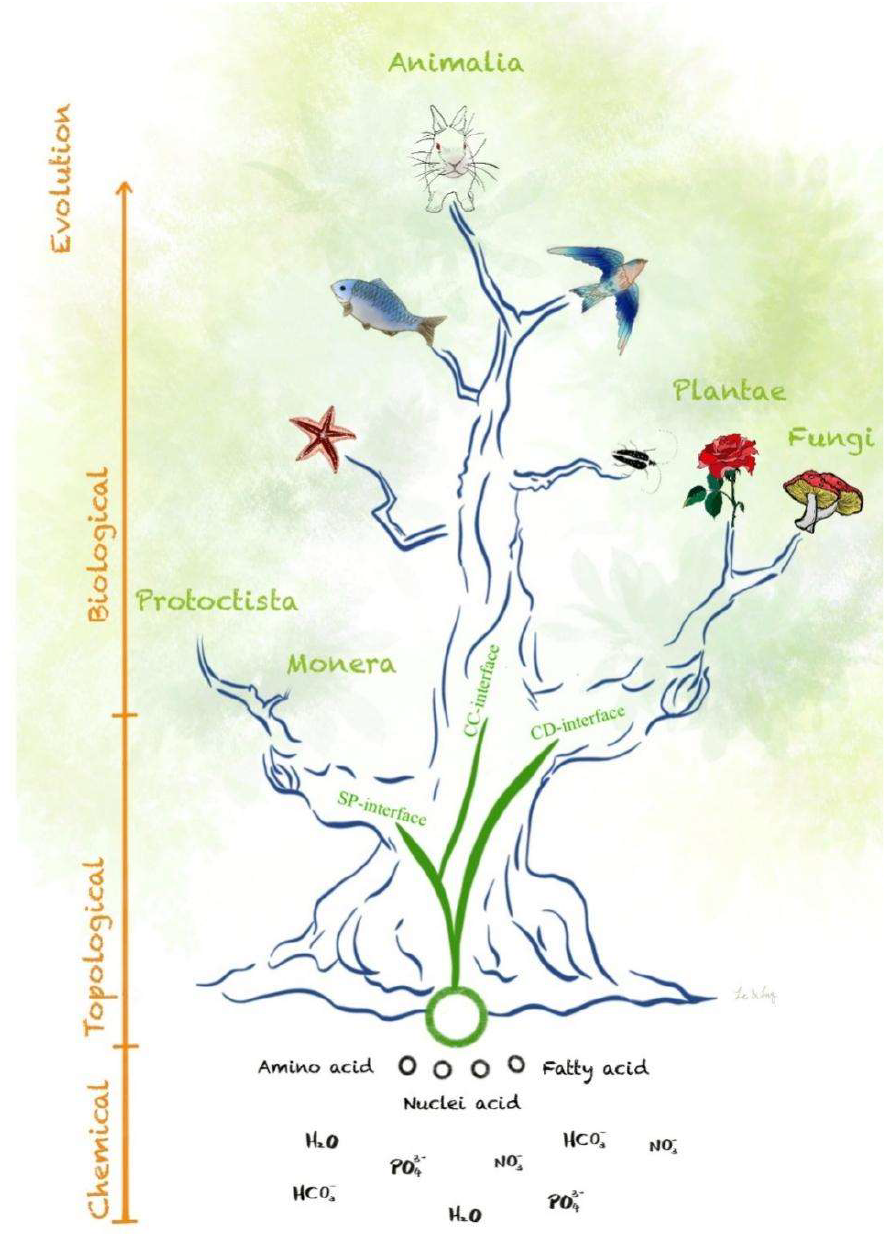
Illustration of “the bud of life” and topological evolution. Topological evolution is proposed as an intermediate stage between chemical and biological evolution. **Notes:** Black small circles, micelles; large green circle, unicellular organism. SP-, CD-, and CC-interface refer to as spherical-, closed disk-, and closed-cylinder-interface between organismal systems and environment, respectively.

## Abbreviations

CC: closed cylinder
CD: closed disk
GA: geometrically allowed
HF: homeostatically favored
NRT: non-resource transport
SP: spherical surface
TF: transitional form.

## Acknowledgments

We thank Dr. Jianqiang Yang for comments on topological analysis, Mr. Bo Gao for suggestions on statistical analysis, Mr. Weitao Li and Miss Chen Li for drawing some figures, Drs. Stuart Newman, Hugh Pritchard, Zhe-Kun Zhou, and Xuemin Wang for their suggestion and comments, Drs. Mulan Wang and Miss Jing Ling for critical reading of the manuscript.

## Author contributions

Conceptualization, WQL. Methodology, WQL and XDZ. Modeling and simulation, WQL. Investigation, XDZ. Funding acquisition, WQL and XDZ. Project administration, WQL. Supervision, WQL. Writing – original draft, WQL. Writing – review & editing, WQL and XDZ.

## Data and materials availability

All data are available in the main text or the supplementary materials.

## Supplementary Materials

Methods

Tables S1–S6.

Figs. S1–S3.

Data S1–S4.

