## Supplementary Materials for "Organism-Environment Topological Interfaces Drive the Origination of Organismal Form"

**This PDF file includes:**

Methods.

Tables S1–S6.

Figs. S1–S3.

### Methods

#### 1. Formula for stoichiometric homeostasis in complex systems

The mathematical relation between environmental stoichiometry ($X_0$) and systemic stoichiometry ($X_1$) (Fig. 1b) can be expressed as Eq. (1).

$$X_1 = aX_0^b \quad (1)$$

where $a$ is a constant. For simplicity, we define $a = 1$ hereafter, yielding Eq. (1-1).

$$X_1 = X_0^b \quad (1\text{-}1)$$

We posit a complex system comprising subsystems 1, 2, …, $i$, where resources flow sequentially from the environment ($X_0$), into the subsystems ($X_1$, $X_2$, …, $X_i$) under homeostatic regulation by each subsystem ($b_1$, $b_2$, …, $b_i$; $0 \le b_i \le 1$). Based on Eq. (1-1), we derive Eqs. (1-2) – (1-4)..

$$X_1 = X_0^{b_1} \quad (1\text{-}2)$$

$$X_2 = X_1^{b_2} = (X_0^{b_1})^{b_2} = X_0^{b_1 b_2} \quad (1\text{-}3)$$

$$X_i = X_{i-1}^{b_i} = \cdots = X_0^{b_1 b_2 \cdots b_i} \quad (1\text{-}4)$$

Let the biomass of subsystem $i$ be $M_i$ (moles). The molar fraction of subsystem $i$, denoted $F_i$, is defined by Eq. (1-5).

$$F_i = \frac{M_i}{M_1 + M_2 + \cdots + M_i} \quad (1\text{-}5)$$

Assuming that subsystemic biomass is proportional to its volume, we calculate $F_i$ from the volume $V_i$ using Eq. (1-6).

$$F_i = \frac{V_i}{V_1 + V_2 + \cdots + V_i} \quad (1\text{-}6)$$

The total stoichiometry of the system is then obtained via Eq. (1-7).

$$X_{\text{total}} = \sum_i F_i \times X_i \quad (1\text{-}7)$$

Substituting Eq. (1-4) into Eq. (1-7) gives Eq. (1-8).

$$X_{\text{total}} = \sum_i F_i \times X_0^{b_1 b_2 \cdots b_i} \quad (1\text{-}8)$$

### 2. Formulation of systemic stoichiometry under fluctuating environmental stoichiometry

Given that environmental stoichiometry $X_0 > 0$, we assume its fluctuation follows a sine function of time, defined by Eq. (2-0).

$$X_0 = A(\text{Sin}t + 1) \quad (2\text{-}0)$$

where $t$ is time and $A$ is a constant representing the maximum and minimum of $X_0$. For simplicity, we define $A = 1$, yielding Eq. (2).

$$X_0 = (\text{Sin}t + 1) \quad (2)$$

Combining Eq. (1-1) and (2), we obtain Eq. (3).

$$X_1 = (\text{Sin}t + 1)^b \quad (3)$$

Combining Eq. (1-8) and (2), we derive the total stoichiometry for a system containing $i$ subsystems, defined by Eq. (3-1).

$$X_{\text{total}} = \sum_i F_i \times (\text{Sin}t + 1)^{b_1 b_2 \cdots b_i} \quad (3\text{-}1)$$

### 3. Formulation of stoichiometric variation in the closed-disk-interface system with increasing complexity

We constructed three closed-disk (CD)-interface systems comprising single, double, and triple subsystems (Fig. 2a–c). The CD-interface system with a single subsystem

(Fig. 2a) has the mole fraction $F_1$ = 1. Based on Eq. (3-1), we obtain systemic stoichiometry ($X_{\mathrm{S}}$) defined by Eq. (5).

$$X_{\mathrm{S}} = (\mathrm{Sin}t + 1)^{b_1} \quad (5)$$

For a CD-interface system with two subsystems (subsys. 1 and 2; Fig. 2b), we assume equal biomass between subsystems for simplicity, giving $F_1 = F_2 = \frac{1}{2}$. Based on Eq. (3-1), we define systemic stoichiometry via Eq. (6).

$$X_{\mathrm{D}} = \frac{1}{2}[(\mathrm{Sin}t + 1)^{b_1} + (\mathrm{Sin}t + 1)^{b_1 b_2}] \quad (6)$$

For a CD-interface system with three subsystems (subsys. 1, 2, and 3; Fig. 2c), we assume equal biomass among subsystems, giving $F_1 = F_2 = F_3 = \frac{1}{3}$ . Based on Eq. (3-1), we obtain the systemic stoichiometry using Eq. (7).

$$X_{\mathrm{T}} = \frac{1}{3}[(\mathrm{Sin}t + 1)^{b_1} + (\mathrm{Sin}t + 1)^{b_1 b_2} + (\mathrm{Sin}t + 1)^{b_1 b_2 b_3}] \quad (7)$$

### 4. Formulation of stoichiometric variation in spherical-interface system with increasing complexity

We constructed three spherical (SP)-interface systems comprising single, double, and triple subsystems (Fig. 2e–g). Subsystems develop inward along the normal direction of the interface, occupying a small spherical shell or sphere. We assume the system has a radius ($r$) of 1, and that the subsystems have equal thickness along $r$ (i.e., in the normal direction). We calculate the volume of each subsystem and its molar fraction $F_i$ based on Eq. (1-6). The volumes and molar factions of subsystemic $F_i$ are listed in Table S1. Substituting $F_i$ into Eq. (3-1), we obtain systemic stoichiometry via Eqs. (8) – (10).

$$X_{\mathrm{S}} = (\mathrm{Sin}t + 1)^{b_1} \quad (8)$$

$$X_{\mathrm{D}} = 0.875 \times (\mathrm{Sin}t + 1)^{b_1} + 0.125 \times (\mathrm{Sin}t + 1)^{b_1 b_2} \quad (9)$$

$$X_{\mathrm{T}} = 0.704 \times (\mathrm{Sin}t + 1)^{b_1} + 0.259 \times (\mathrm{Sin}t + 1)^{b_1 b_2} + 0.037 \times (\mathrm{Sin}t + 1)^{b_1 b_2 b_3} \quad (10)$$

### 5. Formulation of stoichiometric variation in the closed-cylinder-interface system with increasing complexity

We constructed three closed-cylinder (CC)-interface systems comprising single, double, and triple subsystems (Fig. 2i–k). Subsystems develop outward along the normal direction of the interface, forming larger cylindrical shell. We assume the base radius (*r*) of the systemic cylindrical shell is 1, and the cylindrical shells of subsystems have equal thickness along *r* (i.e., in the normal direction). We calculate the volume of each subsystem and its molar fraction $F_i$ based on Eq. (1-6). The volumes and molar factions $F_i$ of the subsystems are listed in Table S2. Substituting $F_i$ into Eq. (3-1), we obtain systemic stoichiometry via Eqs. (11) – (13).

$$X_{\mathrm{S}} = (\mathrm{Sin}t + 1)^{b_1} \quad (11)$$

$$X_{\mathrm{D}} = 0.25 \times (\mathrm{Sin}t + 1)^{b_1} + 0.75 \times (\mathrm{Sin}t + 1)^{b_1 b_2} \quad (12)$$

$$X_{\mathrm{T}} = 0.111 \times (\mathrm{Sin}t + 1)^{b_1} + 0.333 \times (\mathrm{Sin}t + 1)^{b_1 b_2} + 0.556 \times (\mathrm{Sin}t + 1)^{b_1 b_2 b_3} \quad (13)$$

### 6. Formulation of stoichiometric variation in closed-cylinder-interface system with parallel stoichiometric gradients

We constructed three systems comprising single, double, and triple subsystems arranged in parallel along the closed-cylinder (CC)-interface representing environmental stoichiometric flux of zero, one, and two gradients within the CC-interface, respectively (Fig. 4b–d).

For the system with a single subsystem (Fig. 4b), the mathematical relationship between subsystemic stoichiometry ($Y_1$) and environmental stoichiometry ($X_0$) is $Y_1 = X_0^{b_1} = (\mathrm{Sin}t + 1)^{b_1}$(Eqs. 1 and 2-1). The systemic stoichiometry is then be expressed via Eq. (14).

$$Y_{\mathrm{S}} = Y_1 = X_0^{b_1} = (\mathrm{Sin}t + 1)^{b_1} \quad (14)$$

For the system with two subsystems (Fig. 4c), environmental stoichiometric flux creates a gradient from entrance $X_0$ to exit $X_1$ along the CC interface, with a gradient coefficient of $f=\frac{X_1}{X_0},(0<f\leq 1)$. The corresponding stoichiometries of $X_0$ and $X_1$ in the subsystems are $Y_1$ and $Y_2$, respectively (Fig. 5c). We obtain $Y_1=X_0^{b_1}$ and $Y_2=X_1^{b_2}$ from Eq. (1). Substituting in $X_0=(\mathrm{Sin}t+1)$ (Eq. 2-1) and $X_1=fX_0$, the two subsystemic stoichiometries are expressed as $Y_1=(\mathrm{Sin}t+1)^{b_1}$ and $Y_2=f^{b_2}(\mathrm{Sin}t+1)^{b_2}$, respectively. Assuming equal biomass between subsystems, their molar fractions are $F_1=F_2=\frac{1}{2}$. We then obtain the systemic stoichiometry via Eq. (15).

$$Y_{\mathrm{D}}=\frac{1}{2}Y_1+\frac{1}{2}Y_2=\frac{1}{2}[(\mathrm{Sin}t+1)^{b_1}+f^{b_2}(\mathrm{Sin}t+1)^{b_2}] \quad (15)$$

For the system with three subsystems (Fig. 4d), environmental stoichiometric flux creates two gradients (from $X_0$ to $X_1$ and $X_1$ to exit $X_2$) along the CC-interface. For simplicity, we assume equal gradients with $f=\frac{X_1}{X_0}=\frac{X_2}{X_1},(0<f\leq 1)$ . The corresponding stoichiometries in the subsystems for $X_0$, $X_1$, and $X_2$ are $Y_1$, $Y_2$, and $Y_3$, respectively. We obtain $Y_1=X_0^{b_1}$, $Y_2=X_1^{b_2}$, and $Y_3=X_2^{b_3}$ from Eq. (1). Substituting $X_0=(\mathrm{Sin}t+1)$ (Eq. 2-1), $X_1=fX_0$, and $X_2=fX_1=f^2X_0$, the three subsystemic stoichiometries are expressed as $Y_1=(\mathrm{Sin}t+1)^{b_1}$, $Y_2=f^{b_2}(\mathrm{Sin}t+1)^{b_2}$, and $Y_3=f^{2b_3}(\mathrm{Sin}t+1)^{b_3}$, respectively. Assuming equal biomass among subsystems, their molar fraction are $F_1=F_2=F_3=\frac{1}{3}$. We then obtain systemic stoichiometry via Eq. (16).

$$Y_{\mathrm{T}}=\frac{1}{3}Y_1+\frac{1}{3}Y_2+\frac{1}{3}Y_3=\frac{1}{3}[(\mathrm{Sin}t+1)^{b_1}+f^{b_2}(\mathrm{Sin}t+1)^{b_2}+f^{2b_3}(\mathrm{Sin}t+1)^{b_3} \quad (16)$$

**Table S1. Calculated molar fractions of spherical (SP)-interface systems comprising single, double, or triple subsystems (Fig. 2e–g)).** The SP-interface system has a radius ($r$) of 1, and subsystems are equal thickness along $r$ (i.e., in the normal direction). The volume of each subsystem can be calculated using the formula $\frac{4}{3}\pi r^3$, and the molar fraction ($F_i$) is derived from Eqs. (1-6).

| **System** | **Subsystem No.** | **Radius** | **Volume (π)** | **Molar fraction ($F_i$)** |
|---|---|---|---|---|
| Containing single subsystem | 1 | 1 | 4/3 | 1 |
| Containing double subsystem | 1 | 1 | 28/24 | 0.875 |
| | 2 | 1/2 | 4/24 | 0.125 |
| Containing triple subsystem | 1 | 1 | 76/81 | 0.704 |
| | 2 | 2/3 | 28/81 | 0.259 |
| | 3 | 1/3 | 4/81 | 0.037 |

**Table S2. Molar fractions of closed cylinder (CC)-interface systems comprising single, double, or triple subsystems (Fig. 2i–k).** The systemic cylindrical shell has a radius (*r*) of 1, and the cylindrical shells of subsystems have equal thickness along *r* (i.e., in the normal direction). The volume of each subsystem is calculated using $\pi r^2 h$ (where $h$ is the cylinder height), and the molar fraction ($F_i$) of each subsystem is derived from Eqs. (1–6).

| **System** | **Subsystem No.** | **Radius of base** | **Height** | **Volume (π)** | **Molar fraction ($F_i$)** |
|---|---|---|---|---|---|
| Containing single subsystem | 1 | 1 | $h$ | $h$ | 1 |
| Containing double subsystem | 1 | 1/2 | $h$ | $\frac{1}{4}h$ | 0.25 |
| | 2 | 1 | $h$ | $\frac{3}{4}h$ | 0.75 |
| Containing triple subsystem | 1 | 1/3 | $h$ | $\frac{1}{9}h$ | 0.111 |
| | 2 | 2/3 | $h$ | $\frac{3}{9}h$ | 0.333 |
| | 3 | 1 | $h$ | $\frac{5}{9}h$ | 0.556 |

**Table S3. Representation of spherical (SP), closed disk (CD), and closed-cylinder (CC) interfaces for the organismal forms at the kingdom level.** Resource intake characteristics: SP, resource intake from the outside of the interface; CD, resource intake from one side of the interface; CC, resource intake from the inside of the interface. “ √ ” indicates that the primary feature of the indicated kingdom matches the topological interface; “–” indicates not applicable.

| **Broad classification system** | | **Observed primary characteristics for resource intake** [1,2] | **Can be represented by topological interface** | | |
|---|---|---|---|---|---|
| | | | SP | CD | CC |
| Two-kingdom | *Plantae* | Nutrient absorption by roots | – | √ | |
| | *Animalia* | Food ingestion through cavities | | | √ |
| Three-kingdom | *Protoctista* | Nutrient absorption + Unicellular | √ | | |
| | *Plantae* | Nutrient absorption + Multicellular | | √ | |
| | *Animalia* | Food ingestion + Multicellular | | | √ |
| Four-kingdom | *Monera* | Nutrient absorption + Unicellular + Prokaryotic | √ | | |
| | *Protoctista* | Nutrient absorption + Unicellular + Eukaryotic | √ | | |
| | *Plantae* | Nutrient absorption + Multicellular + Eukaryotic | | √ | |
| | *Animalia* | Food ingestion + Multicellular + Eukaryotic | | | √ |
| Five-kingdom | *Monera* | Nutrient absorption + Unicellular + Prokaryotic | √ | | |
| | *Protista* | Nutrient absorption + Unicellular + Eukaryotic | √ | | |
| | *Plantae* | Nutrient absorption + Multicellular + Eukaryotic + Photosynthesis | | √ | |
| | *Fungi* | Nutrient absorption + Multicellular + Eukaryotic + No photosynthesis | | √ | |
| | *Animalia* | Food ingestion + Eukaryotic + Multicellular | | | √ |

[1]R. H. Whittaker, New concepts of kingdoms or organisms. Evolutionary relations are better represented by new classifications than by the traditional two kingdoms. Science 163, 150-160 (1969).

[2]R. H. Whittaker, On the broad classification of organisms. The Quarterly review of biology 34, 210-226 (1959).

153

**Table S4. Chi-square tests (2×2 contingency tables, *df* = 1) for four independent binary morphological features**

The observed and predicted phylum numbers are from Data S2, S2, S3, and S4, respectively. The results showed extremely significant correlations between observed and predicted types.

| **1. Size-related feature** ($n = 52$, $\chi^2 = 36.903$, $p < 0.000001$) | | |
|---|---|---|
| Prediction→ Observation↓ | With spatial constraint (small) | Without spatial constraint (large) |
| Microscopic | 20 | 2 |
| Macroscopic | 2 | 28 |

| **2. Complexity-related feature** ($n = 52$, $\chi^2 = 44.571$, $p < 0.000001$) | | |
|---|---|---|
| Prediction→ Observation↓ | Not favoring subsystem development (simple) | Favoring subsystem development (complex) |
| Without tissue | 24 | 2 |
| With tissues | 0 | 26 |

| **3. Motility-related feature** ($n = 52$, $\chi^2 = 37.284$, $p < 0.000001$) | | |
|---|---|---|
| Prediction→ Observation↓ | Parallel-stoichiometric gradient | No Parallel-stoichiometric gradient |
| Motile | 31 | 4 |
| Nonmotile | 0 | 17 |

| **4*. Animal size-related feature** ($n = 28$, $\chi^2 = 16.121$, $p = 0.000059$) | | |
|---|---|---|
| Prediction→ Observation↓ | Less space for subsystem development | More space for subsystem development |
| Small | 6 | 0 |
| Large | 3 | 19 |

* Fisher's Exact Test, $p = 0.000223 < 0.01$.

**Table S5. Predicted and observed species proportions at the kingdom level.** SP, spherical; CD, closed disk; CC, closed cylinder. Numbers in parentheses represent observed species counts.

| **Item** | **Percentage of species number** | | | **Note and reference** |
|---|---|---|---|---|
| Topological model | SP-interface organism | CD-interface organism | CC-interface organism | |
| Kingdom | *Monera* & *Protista* | *Plantae* & *Fungi* | *Animalia* | See Table S3 |
| Predicted | 4% | 24% | 72% | See Fig. S4 |
| Observed | 5% | 22% | 70-73% | Science **241**:1441(1998) |
| | 1.9% (27749) | 19.9% (286504) | 78.2% (1124516) | PLoS Biol **9**: e1001127 (2011) |
| | 0.3% （5437） | 29. 7% (483702) | 70.0% (1140884) | Species2000 (https://www.sp2000.org) |

**Table S6. Explanation for Cambrian Explosion features.** SP-, CD-, and CC-interface systems correspond to microbes (*Monera* and *Protista*), plants (*Plantae* and *Fungi*), and animals (Animalia) at the kingdom level, respectively (Section 3.1; Fig. 2; Table S3).

| Feature of the Cambrian Explosion | | Can the feature be explained? | | |
|---|---|---|---|---|
| | | **By Darwinian Theory** | **By Modern Synthesis** | **By Topological Interface (This study)** |
| 1 | **Temporal concentration** (Major animal groups appeared within an extremely short geological time span, approximately 10–20 million years) | **No** (The theory emphasizes gradual evolution, making it difficult to explain such a sudden, concentrated burst in a short geological time span) | **Partially** (The theory allows for rapid adaptive radiations, but still struggles to explain such an extremely tightly clustered event in geological time span) | **Yes** (Compared with SP- and CD-interface systems, CC-interface system has a stronger homeostatic tendency and quadratically larger space to develop subsystems, allowing it to mature far faster. Thus, a rapid burst of animals has occurred) |
| 2 | **Morphological complexity leap** (Involved the sudden emergence of animals with complex body structures, e.g., organs, body segments, symmetry) | **No** (The theory lacks an explanation for the rapid emergence of complex body structures, due to limited understanding of genetics and development) | **Partially** (The theory can explain some morphological innovations via "genetic variation + natural selection", but it does not explain why the complexity leaps) | **Yes** (Compared with SP- and CD-interface systems, CC-interface system has a stronger homeostatic tendency and quadratically expanding space to increase subsystem numbers, allowing its complexity to leap. Thus, animal structures become dramatically more complex) |
| 3 | **Rapid diversification** (A large number of new animal groups and morphotypes appeared in a very short time, leading to a sharp increase in biodiversity) | **No** (The theory assumes slow accumulation of small variations, and thus struggles to explain rapid appearance of many new groups). | **Partially** (The theory can account for how animal life diversifies via natural selection, genetic variation, and ecological opportunity, but it does not explain why this diversification occurs so rapidly) | **Yes** (Compared with SP- and CD-interface systems, CC-interface system develops subsystems the fastest and increases their numbers most rapidly. Thus, many new animal morphotypes appeared in a very short time) |
| 4 | **Occurring only in the animal kingdom** (Occurred exclusively in animals, with no comparable event in plants or microbes) | **No** (The theory does not address why this burst of diversification was limited to animals and did not occur in plants or microbes) | **No** (The theory explains increasing animal complexity, but not why this does not apply to plants or microbes) | **Yes** (Compared with CC-interface system, SP-interface system has far less tendency and space for subsystem development, while CD-interface system has less tendency and only linearly expanding space for subsystem development, allowing their subsystems develop gradually and slowly. Thus, plants and microbes diversified incrementally and continuously) |
| 5 | **Occurring only once** (The only major event in Earth's history where so many new animal phyla emerged in a concentrated manner) | **No** (The theory provides no mechanismfor why such a major evolutionary event happened only once in Earth's history) | **No** (The theory lacks a clear explanation for the uniqueness of the Cambrian Explosion as a one-time event) | **Yes** (SP-, CD-, and CC-interface systems appeared sequentially. The latter, with a strong tendency and quadratic expansion to develop subsystems, triggers a subsystem burst, with no secondary bursts nor higher intensity bursts from others. Thus, a burst event like the Cambrian Explosion has occurred only once) |

**Fig. S1. Simulated systemic stoichiometry $Y_D$ (Eq. [15], Fig. 4c) in in closed cylinder (CC)-interface systems under different homeostatic regulation conditions.**

**Notes:** Purple lines, $Y_D$ of systems with subsystemic $b_1 > b_2$; blue lines, $Y_D$ of systems with subsystemic $b_1 = b_2$; green lines, $Y_D$ of systems with subsystemic $b_1 < b_2$; black dotted lines, environmental stoichiometry (equivalent to that of systems with homeostatic regulation constant $b = 1$); red dotted lines, stoichiometry of systems with homeostatic regulation constants $b_i = 0$ ($i = 1$ or 2); $t$, time; $b_1$ and $b_2$, homeostatic regulation constants of subsystem 1 and 2, respectively; $f = \frac{X_1}{X_0}$ $(0 < f \leq 1)$, gradient coefficient.

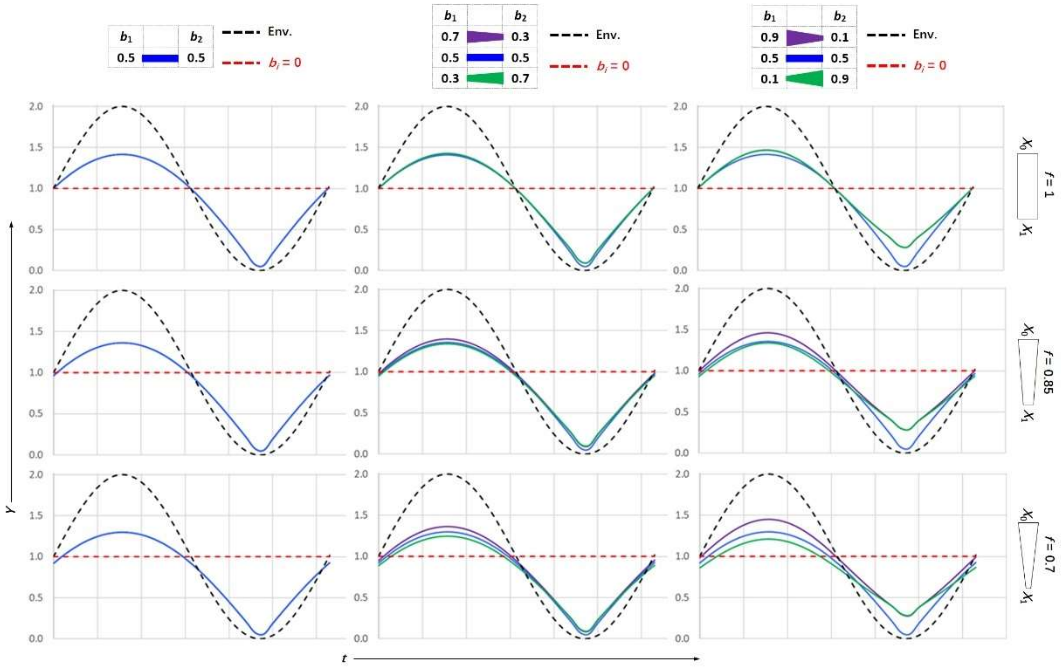

**Fig. S2. Simulated systemic stoichiometry $Y_T$ (Eq. [16]) with two environmental stoichiometry gradients (Fig. 5d) in the CC-interface systems under different homeostatic regulation conditions. Notes:** $Y$, systemic stoichiometry; $t$, time; purple lines, $Y_T$ of systems with subsystemic $b_1 > b_2 > b_3$; blue lines, $Y_T$ of systems with subsystemic $b_1 = b_2 = b_3$; green lines, $Y_D$ of systems with subsystemic $b_1 < b_2 < b_3$; red dotted lines, stoichiometry of systems with $b_i = 0$ ($i$ = 1, 2, or 3); $t$, time; $b_1$, $b_2$, and $b_3$, homeostatic regulation constants of subsystem 1, 2, and 3, respectively (Methods 3–5); $f = \frac{X_1}{X_0} = \frac{X_2}{X_1}$ $(0 < f \leq 1)$, gradient coefficient.

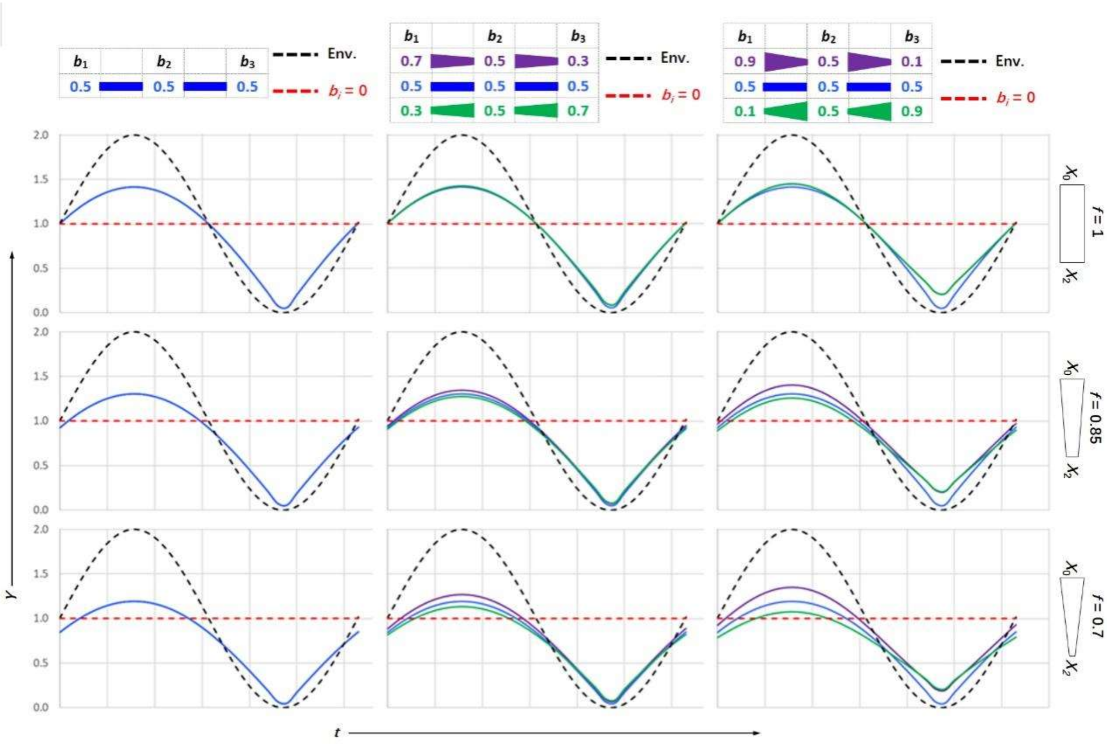

**Fig. S3. Morphological space diagrams, calculations of space ratios for newly developed subsystems, and calculations of species proportion at kingdom level. Notes:** Blue area, environment; red area, system; yellow area, newly developed subsystem; boundary between red and blue areas, topological interface between environment and system; black arrows, normal direction of interfaces for subsystem development; *r*, radius of sphere or cylinder base; *h*, cylinder height.

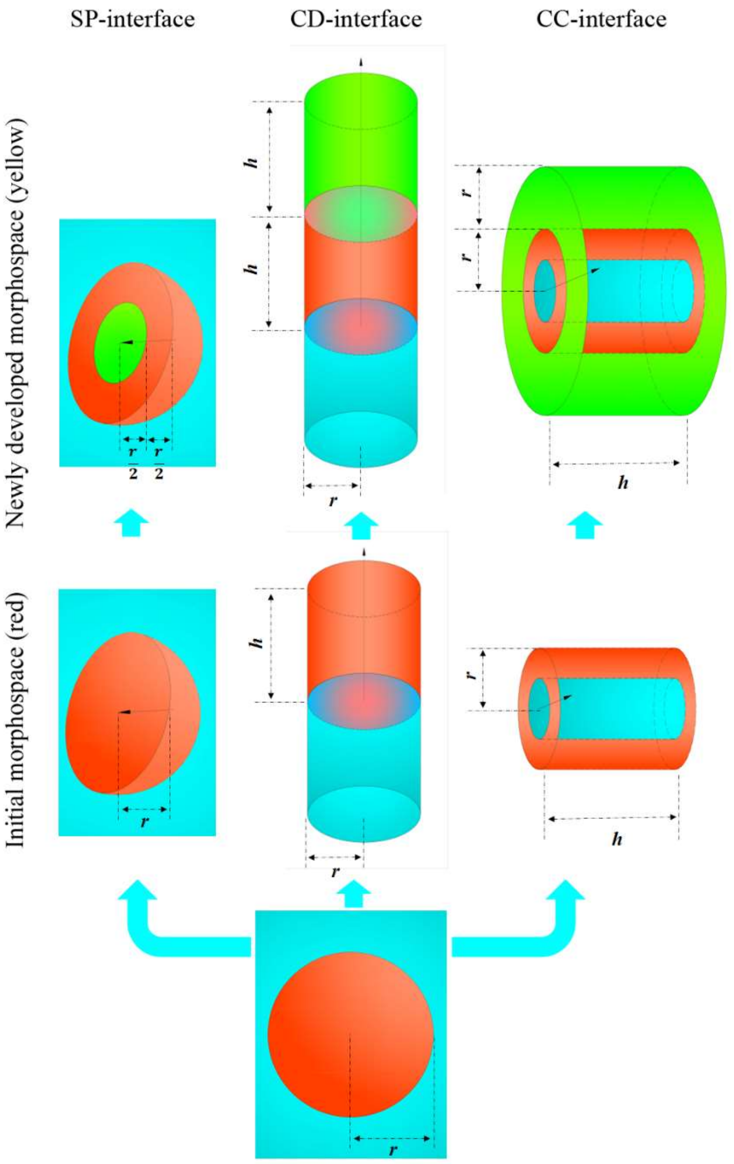


**Calculations of space ratios for newly developed subsystems:** Morphospace volumes of newly developed subsystems are calculated based on our proposed mechanism (Fig. 3). For comparative consistency, all systems are assumed to have equal initial size ($r = 1$, $h = 1$) and to develop new subsystems at a 1:1 length ratio along the interface normal directions.

SP-interface organisms (left panel): Initial volume = $\frac{4}{3}\pi r^3 = \frac{4}{3}\pi$. The new subsystems develop inward from the interface with $r = \frac{1}{2}$ (the 1:1 ratio bisects the initial normal dimension). Volume of newly developed subsystems: $V_{\text{SP}} = \frac{4}{3}\pi(\frac{1}{2})^3 = \frac{1}{6}\pi$.

CD-interface organisms (middle panel): Initial volume = $\pi r^2 h = \pi$. New subsystems develop onto one side of the interface with $h = 1$ (the 1:1 ratio extends the initial normal dimension). Volume of newly developed subsystems: $V_{\text{CD}} = \pi r^2 h = \pi$.

CC-interface organisms (right panel): Initial volume = $\pi r^2 h = \pi$. New subsystems develop outward from cylindrical interface with $r = 2$ (the 1:1 ratio extends the initial normal dimension). Volume of the newly developed subsystems: $V_{\text{CC}} = \pi(2r)^2 h - \pi r^2 h = 3\pi$.

The space ratios for newly developed subsystems are

$$V_{\text{SP}}:V_{\text{CD}}:V_{\text{CC}} = \frac{1}{6}\pi:\pi:3\pi = \frac{1}{6}:1:3 = 4.0\%:24.0\%:72.0\%$$

**Calculations of species proportion at the kingdom level:** Assuming that SP-, CD-, and CC-interface organisms develop subsystems to generate species diversity, and that morphospace volume (*V*) of new subsystems scales with species number (N), we derive below relationship. Given that SP, CD, and CC interfaces represent *Monera* + *Protoctista*, *Plantae* + *Fungi*, and *Animalia*, respectively (Table S3), the species proportion of these kingdoms is defined as:

$$N_{\text{SP}}:N_{\text{CD}}:N_{\text{CC}} = V_{\text{SP}}:V_{\text{CD}}:V_{\text{CC}} = \frac{1}{6}:1:3 = 4.0\%:24.0\%:72.0\%$$

**Data S1. (Separate .xlsx file)**

Data S1. Representation of topological interfaces for organismal forms at the phylum level

**Data S2. (Separate .xlsx file)**

Data S2. Comparison of observed and predicted primary morphological features

**Data S3. (Separate .xlsx file)**

Data S4. Comparison of observed and predicted body sizes in animals

**Data S4. (Separate .xlsx file)**

Data S4. Comparison of observed and predicted body sizes in animals
